# Differential impacts of various bacteria on the health and survival of the sea anemone *Exaiptasia diaphana*

**DOI:** 10.1101/2025.02.13.638178

**Authors:** Kaitlyn M. Romo, Cawa Tran

## Abstract

Coral reefs are threatened by rising ocean temperatures associated with climate change, leading to a breakdown of the cnidarian-dinoflagellate symbiosis and growth of pathogenic bacteria. The host response to diverse bacteria can vary from species to species. In this study, the sea anemone *Exaiptasia diaphana* (commonly referred to as ‘Aiptasia’), a prominent laboratory model for coral-symbiosis research, was used to investigate host responses to three key bacterial species associated with both Aiptasia and corals. *Ruegeria mobilis*, *Vibrio alginolyticus*, and *Alteromonas macleodii* were individually inoculated into the seawater of both symbiotic (with dinoflagellates) and aposymbiotic (without dinoflagellates) hosts to determine their effects on host health. Each bacterium was hypothesized to have a different effect on host health— beneficial, pathogenic, or no effect. Various densities of each bacterium were used to assess impacts on host morphology and survival over 14 days at 27°C. Host biomass, protein content, and dinoflagellate abundance (to assess bacterial effects on host bleaching) were examined over 7 days at 27°C in response to an inoculation density of 8×10^8^ cells ml^-1^ of each bacterium. When compared to un-inoculated Aiptasia, *V. alginolyticus* reduced host survival of symbiotic and aposymbiotic hosts, and biomass by 27% and 54% in symbiotic and aposymbiotic Aiptasia, respectively. *V. alginolyticus* also diminished dinoflagellate abundance by 20% in symbiotic Aiptasia. In contrast, *R. mobilis* enhanced survival of symbiotic Aiptasia, biomass by 9%, protein content by 50%, and dinoflagellate abundance by 60%, but had no impacts on aposymbiotic Aiptasia. *A. macleodii* had no major effects on host health at all. As a result, *V. alginolyticus* generally has negative impacts on host health, while *R. mobilis* appears to be a beneficial bacterium. Moreover, host susceptibility to bacteria is dependent on whether animals have their dinoflagellates to begin with and, therefore, bleached cnidarians may experience more drastic responses to pathogens like *V. alginolyticus*, but no benefits from a potential probiotic like *R. mobilis*.

**Importance:** Identification of specific beneficial microbes is a major contribution to conservation techniques, such as microbiome manipulation, that aim to assist cnidarians in adapting to climate change. However, to maintain these conservation efforts in the long term, moving forward, we must prioritize foundational questions that examine dinoflagellate-bacterial interactions and bacterial-colonization dynamics within the cnidarian host.

## Introduction

Coral health is dependent upon the microbial symbioses that corals have to maintain with photosynthetic dinoflagellates and various types of prokaryotes. As ocean temperatures continue to rise, coral-reef declines are expected to occur more frequently (Hoegh-Guldberg et al. 1999) when these symbiotic relationships become compromised. Corals expel the algae and lose photosynthetic abilities along with their coloration, in a phenomenon known as coral bleaching (Yellowlees et al. 2008). At the same time, elevated temperatures can drive the growth of opportunistic and pathogenic bacteria, production and release various virulence factors, which can lead to disease within the host (Rosenberg et al. 2004; Littman et al. 2010).

Corals have extremely diverse and complex microbiomes which they may rely on for essential functions, though the present bacterial community is subjected to constant environmental fluctuations. Corals may depend on bacteria to perform specific activities that include protection from pathogens (Ritchie 2006; Krediet et al. 2013; Glasl et al. 2016; Miura et al. 2019), cycling of various nutrients (C, N, S) for energy sources (Rädecker et al. 2015; Peixoto et al. 2017), and acclimation to heat stress (Rosado et al. 2019). Changes in the environmental bacterial community, therefore, present a challenge for the establishment of a suitable microbiome. The establishment and maintenance of key bacteria in the coral microbiome must be specific, yet there is no consensus on how bacteria are selected by corals (Krediet et al. 2013; Leite 2017).

Corals obtain their microbiome through vertical (parent to offspring) or horizontal (from the environment) transmission, or a combination of both (Leite et al. 2017; Damjanovic et al. 2020). Vertical transmission provides and maintains mutualistic partners, while horizontal transmission provides the benefit of acquiring symbionts adapted to a specific environment (Byler et al. 2013), often controlled through selection by coral mucus (Ritchie 2006; Koehler et al. 2018). Coral mucus plays an important role as a first defense against pathogenic bacteria (Rivera-Ortega and Thomé 2018), yet this function may also be compromised by rising ocean temperatures. Mucus obtained from corals in naturally elevated temperatures did not inhibit invasive bacteria, suggesting that an increase in ocean temperatures can impact the coral’s natural ability to select against invasive species (Ritchie 2006). This may be a result of a decline in beneficial bacterial species that are typically present to outcompete pathogenic bacteria, such as *Vibrio coralliilyticus* and *Serratia marcescens* (Glasl et al. 2016).

As studies continue to ascertain the role of bacteria within the coral holobiont, microbiome manipulation (Blackall et al.) has been proposed as a possible method to introduce beneficial bacteria into corals to assist them in acclimation to heat stress (Peixoto et al. 2017, Rosado et al. 2019). Putatively beneficial microorganisms for corals (pBMC’s) with key traits— including the ability to catabolize dimethylsulfoniopropionate, fix nitrogen, and/or inhibit pathogens—were identified and inoculated into corals *in vitro*, inducing them to become less susceptible to bleaching at elevated temperatures (Rosado et al. 2019). Though microbiome manipulation as a novel approach appears to be a promising coral-conservation effort, bacteria have various functions and may influence microbiome structure and integrity in more ways than one. Thus, there is still much to be known about various and combinatorial effects of these bacteria on host health.

The sea anemone *Exaiptasia diaphana* (hereafter ‘Aiptasia’) is commonly employed as a model system for coral research, given their symbiosis with Symbiodiniaceae and some common marine bacteria (Weis et al. 2008; Röthig et al. 2016; Damjanovic et al. 2019). Unlike corals, Aiptasia can be grown with their algal symbionts (symbiotic) or without their algal symbionts (aposymbiotic), making this model ideal for studying symbiosis and bleaching under laboratory conditions (Weis et al. 2008). Aiptasia asexually reproduces rapidly, making propagation of genetically distinct clonal lines fast and easy (Weis et al. 2008; Sunagawa et al. 2009). Only recently, Aiptasia has been used as a model system for microbiome studies. Like corals, the Aiptasia microbiome seems to be dependent upon environmental conditions and algal presence, with bacterial partners changing as temperatures change and as bleaching occurs (Ainsworth and Hoegh-Guldberg 2009; Röthig et al. 2017; Ziegler et al. 2017; Sydnor et al. 2023). Aiptasia is a good model for studying bacterial diseases within corals. When Aiptasia was infected with two well-known coral pathogens, *Vibrio coralliilyticus* and *Vibrio shiloi*, disease progressed comparably to that in corals (Zaragoza et al. 2014). Finally, lab-maintained Aiptasia has a comparable microbiome to Aiptasia that grow natively on the Great Barrier Reef, suggesting that studies done in the lab apply to the wild (Hartman et al. 2020).

In the present study, we used Aiptasia to investigate three bacterial species known to be highly associated with both Aiptasia and corals, and abundant in marine environments (Binsarhan 2015; Hernandez-Agreda et al. 2018; Huggett and Apprill 2018; Sydnor et al. 2023). These bacterial species include a putatively beneficial bacterium (*Ruegeria mobilis*), a putatively pathogenic bacterium (*Vibrio alginolyticus*), and a bacterium that is putatively neither beneficial nor pathogenic (*Alteromonas macleodii*). *R. mobilis* was selected for this study because of its high association with symbiotic and aposymbiotic Aiptasia (Sydnor et al. 2023).

Rhodobacteraceae was the most dominant family in normal and heat-stressed states of Aiptasia, with *Ruegeria* spp. representing a large portion of the Rhodobacteraceae species present (Sydnor et al. 2023). *Ruegeria* spp. is widely associated with corals, abundant in 36 different coral species (Huggett and Apprill 2018). In addition, *R. mobilis* may play an important role in pathogen defense. *Vibrio coralliilyticus* growth was inhibited when *Ruegeria* sp. strains were used in a well-diffusion assay (Miura et al. 2018). *Ruegeria* spp. can metabolize DMSP, a central molecule in the sulfur cycle produced by dinoflagellate algae in coral reefs and other cnidarians. When DMSP is broken down by bacteria, it turns into sulfur-based antimicrobial compounds. These antimicrobial compounds inhibit the growth of coral pathogens, including *V. coralliilyticus* (Peixoto et al. 2017).

*V. alginolyticus* was included in this study because other *Vibrio* spp. cause diseases in corals and Aiptasia (Zaragoza et al. 2014). *Vibrio* was abundantly represented in the Aiptasia microbiome (Binsarhan 2015; Sydnor et al. 2023), and was the most abundant genus in 42 coral species (Huggett and Apprill 2018). In Aiptasia, disease is caused by two well-known coral pathogens, *Vibrio coralliilyticus* and *Vibrio shiloi* (Zaragoza et al. 2014). *V. alginolyticus* is associated with both yellow band disease and *Porites andrewsi* white syndrome in corals (Cervino et al. 2008; Zhenyu et al. 2013). Corals are most susceptible to yellow band disease after their immune system has been compromised by environmental stress factors like rising ocean temperatures. Yellow band disease appears as lesions on the outside of corals and leads to the decimation of entire reefs. White band disease, causing tissue loss, has been connected with various strains of *Vibrio*, which includes *V. alginolyticus* (Zhenyu et al. 2013).

*A. macleodii* was chosen for this study based on its association with Aiptasia and corals (Hernandez-Agreda et al. 2018; Sydnor et al. 2023). *Alteromonas* was dominant in both symbiotic and aposymbiotic Aiptasia, as well as in their seawater (Sydnor et al. 2023). Alteromonadaceae was highly abundant and the most prevalent bacterial family among three coral species from ten reef sites that ranged from 10-80 m below sea level (Hernandez-Agreda et al. 2018). *A. macleodii* has a complex genome, which reflects an adaptable phenotype (López-Pérez et al. 2012). *A. macleodii,* a mesophile and moderate halophile, grows successfully in temperatures that range from 10-40°C, indicating thermotolerance, and can tolerate a wide range (1-18% NaCl) of salinities (López-Pérez et al. 2012).

Diverse parameters were used to assess Aiptasia health in response to bacterial inoculations in this study, including anemone morphology and survival, biomass and protein content (i.e., measurements of stored-resource depletion), and algal abundance (i.e., susceptibility to bleaching). The main goal of this research is to gain an understanding of the impacts key bacteria may play in the health of a cnidarian host, through systematically testing each bacterium to determine whether they have any significant functional roles within the host. These roles may include pertinent interactions with either the cnidarian or its algal symbionts that may change as ocean temperatures change.

## Materials and Methods

### Anemone maintenance and preparation for experiments

To maintain both symbiotic and aposymbiotic anemones of the CC7 clonal line (Sunagawa et al. 2009, Baumgarten et al. 2015), animals were reared in polystyrene tanks that contained 500 ml artificial seawater (ASW) made from Coral Pro Salt (Red Sea, Houston, TX) with a salinity of 32-34 ppt. Clear tanks were used for symbiotic anemones to allow light penetration, and black tanks were used for aposymbiotic anemones to maintain them in the dark. Tanks were kept in an incubator at 27°C on a 12 h light: 12 h dark schedule, with an intensity of 25 μmol photons m^-2^s^-1^ white LED light. Twice a week, anemones were fed freshly hatched *Artemia* nauplii, followed by water changes 8 h later.

Four days before beginning the 7– and 14-d experiments (described below), anemones were primed by placing symbiotic and aposymbiotic animals individually into the wells of 6-well culture plates, with each well containing 6 ml of sterile seawater (SSW), and maintained at 27 °C on the diurnal cycle (described above). The start of the experiment is considered as 0 d. Anemones were not fed and water was not changed for the remainder of each experimental trial.

### Bacterial isolation from Aiptasia and identification by 16S rRNA gene sequence analysis

Bacterial isolates for this project were selected based on traits attributed to each species (**Table 1**). *R. mobilis* and *V. alginolyticus* were previously isolated from the CC7 clonal line of symbiotic Aiptasia, and *A. macleodii* from the H2 symbiotic clonal line (Tran unpublished). Bacteria were isolated by whole-anemone homogenization in SSW using an OMNI Tissue Master (OMNI, Kennesaw, GA), plating serially-diluted homogenates onto marine agar plates (Difco, Detroit, MI), and incubating at 27 ℃ for 48 h. Distinct colonies were selected and streaked onto new marine agar plates to purify and isolate individual bacterial strains, and these plates were incubated at 27 ℃ for 48 h.

**Table 1.**
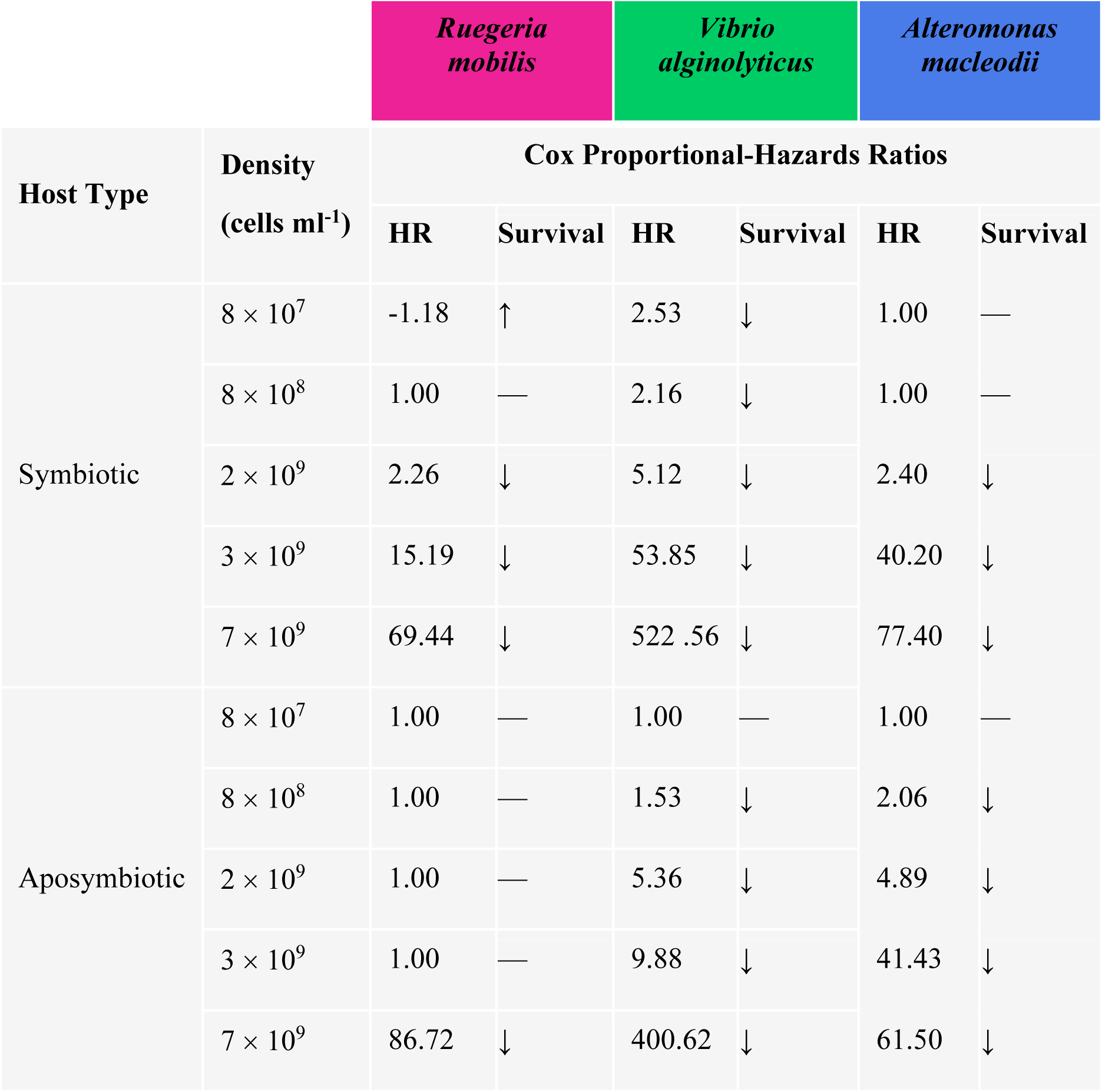
Results of survival and Cox proportional-hazards analysis following inoculation of Aiptasia with three bacterial species, *Ruegeria mobilis*, *Vibrio alginolyticus*, and *Alteromonas macleodii*. Hazards ratios (HR) were determined for each bacterial density in symbiotic and aposymbiotic Aiptasia using the “survival” package in R studio. When HR=1 there was no change in survival; HR<1 there was a decreased hazard; and when HR>1 there was an increased hazard. Change in survival is indicated by arrows, where “↑” indicates increased survival, “↓” indicates decreased survival, and “—” indicates no change.

To identify each bacterial species, the 16S rRNA gene was amplified via PCR using the bacterial-specific primers 8F (5’-AGA GTT TGA TCC TGG CTC AG-3’) and 1492R (5’-GGT TAC CTT GTT ACG ACT T-3’) (Weisburg et al. 1991), with single colonies as DNA templates. The reaction conditions included an initial denaturation at 94°C for 5 min, an annealing temperature of 50°C for 90 s, followed by an amplification phase of 35 cycles at 72°C for 2 min each, and ending at 72°C for 10 min. PCR products were cleaned with the ExoSAP-IT™ PCR Product Cleanup Reagent (Thermo Fisher Scientific, Waltham, MA) according to the manufacturer’s instructions and sent immediately to Elim Biopharmaceuticals, Inc. (Hayward, CA) for sequencing. Nearest relatives were obtained by BLAST searches in the GenBank database on the National Center for Biotechnology Information (NCBI) website. Bacterial species were identified by their closest match with a criterion of >97% similarity. All bacterial isolates have been maintained as 80% glycerol stocks at –80°C.

### Assessing changes in anemone morphology and survival in response to bacterial inoculations

To assess bacterial effects on host survival, 30 symbiotic and 30 aposymbiotic anemones were placed in separate wells of 6-well plates and anemones were primed for 4 d as described above. Overnight cultures of *R. mobilis*, *A. macleodii*, and *V. alginolyticus* were grown in marine broth (Difco, Detroit, MI) for 18 h, shaking at 27°C. The optical density (OD) for each culture was read at 600 nm using a Synergy H1 spectrophotometer microplate reader (BioTek, Winooski, VT) to determine initial bacterial cell concentrations. To prepare for inoculations with Aiptasia, bacterial-cell concentrations were adjusted by replacing the marine broth with the appropriate volumes of SSW to obtain desired inoculation densities. Bacteria were inoculated into wells to achieve the final densities of 0, 8 × 10^7^, 8 × 10^8^, 2 × 10^9^, 3 × 10^9^ and 7 × 10^9^ cells ml^-1^, with five replicate animals for each density (**Figure 1**). Uninoculated anemones maintained under the same conditions served as controls. Following inoculation, anemones were observed for changes in morphology for signs of decreased health (tissue darkening, tentacle retraction, and tissue degradation, as noted in Krediet et al. 2014 and Zaragoza et al. 2014) and survival at 0, 3, 5, 7, 10 and 14 d. Anemones were considered dead if complete tissue degradation occurred. Anemones were photographed with a Leica M165FC stereo microscope with a Leica MC190 camera (7.50 ms exposure, 6.8X gain, 3.4X magnification). The experiment was independently repeated three times, each time with a sample size of 5 anemones per time point for each treatment.

**Figure 1.**
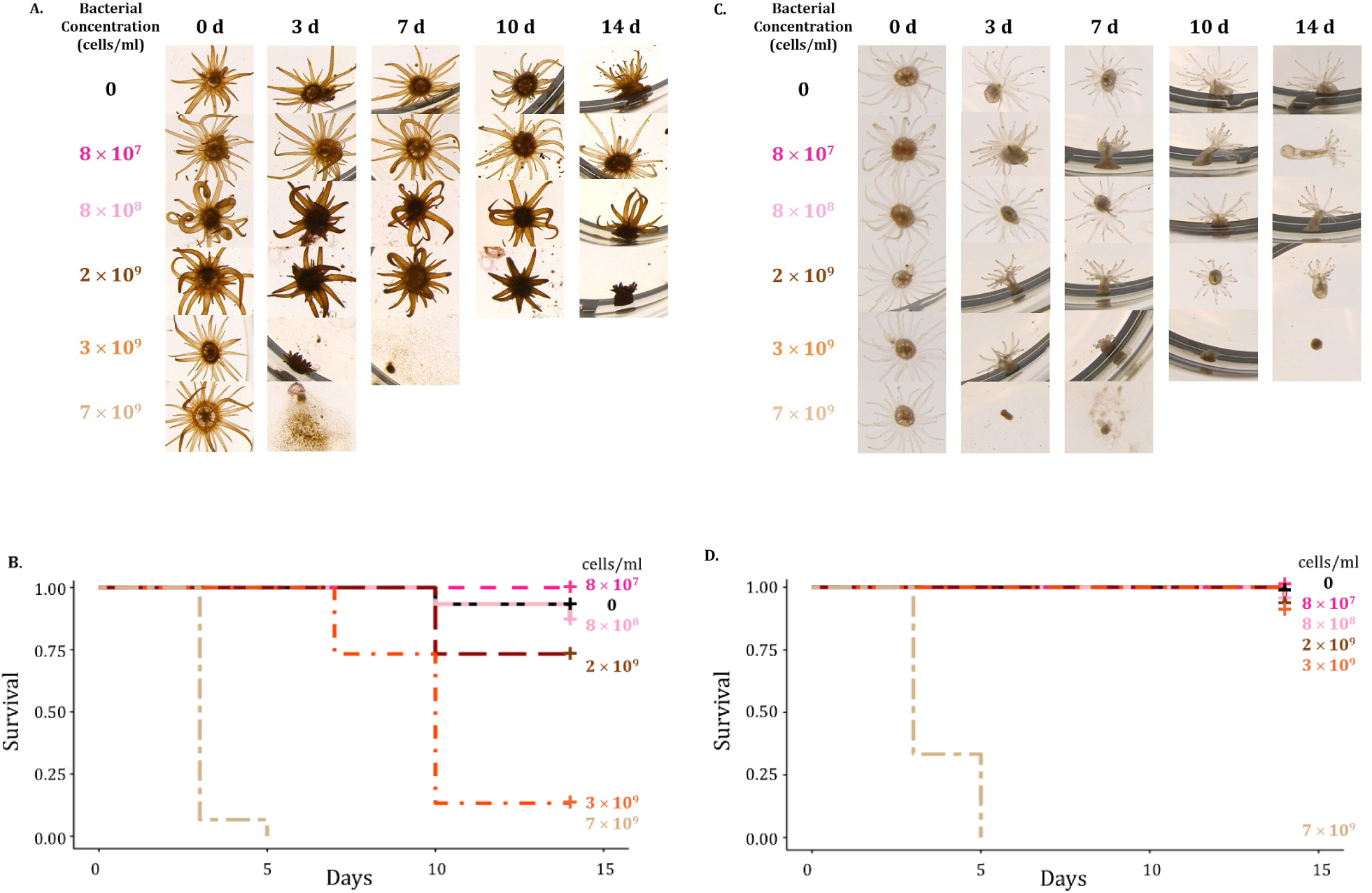
Survival of symbiotic and aposymbiotic Aiptasia over 14 d is dependent on the inoculation density of *Ruegeria mobilis.* Anemones were assessed for signs of disease and death over 14 d following exposure to these densities of *R. mobilis*: 0, 8 × 10^7^, 8 × 10^8^, 2 × 10^9^, 3 × 10^9^ and 7 × 10^9^ cells ml^-1^. Five anemones were assessed for each density and the experiment was independently repeated three times. **A.** Morphology of representative symbiotic Aiptasia under brightfield microscopy shows tissue darkening and tentacle retraction following exposure to *R. mobilis* densities 8×10^8^ cells ml^-1^ and above. Tissue degradation began by 3 d with 3 × 10^9^ and 7 × 10^9^ cells ml^-1^. **B.** Kaplan-Meier survival curves (χ^2^= 121; *df*= 5; *P*<.001) for symbiotic Aiptasia. Plus signs (+) at the end of curves represent censoring (see Materials and Methods). Survival increased at 8 × 10^7^ cells ml^-1^ and decreased at 2 × 10^9^ cells ml^-1^ and above. **C.** Morphology of representative aposymbiotic Aiptasia under brightfield microscopy shows tentacle retraction with densities of 3 × 10^9^ cells ml^-1^ and above. Tissue degradation occurred by 14 d at 3 × 10^9^ cells ml^-1^ and 7 d at 7 × 10^9^ cells ml^-1^. **D.** Kaplan-Meier survival curves (χ^2^= 110; *df*= 5; *P*<.001) aposymbiotic Aiptasia. Plus signs (+) at the end of curves represent censoring (see Materials and Methods). Survival decreased at 7 × 10^9^ cells ml^-1^.

### Quantifying impacts of bacterial inoculations on anemone health

To quantify changes in Aiptasia health in response to bacterial inoculation, 60 symbiotic and 60 aposymbiotic anemones were placed in separate wells of 6-well plates and primed for 4 d as described above. Bacteria were inoculated into wells at a final density of 8 × 10^8^ cells ml^-1^ (**Figure 2**). Uninoculated anemones maintained under the same conditions served as controls. Following inoculation, three metrics were used to assess anemone health at 0, 1, 2, 3, 5, and 7 d: (i) biomass, (ii) protein content, and (iii) algal abundance. Each time point had a sample size of 5 anemones. To measure biomass (defined here as total wet weight), each single anemone was placed on a tared weigh boat, excess water was removed with a pipette, and then weighed on a XE-310 balance (Denver Instrument, Bohemia, NY) in triplicate, with the mass averaged from the three technical replicates. After biomass was recorded, whole anemones were frozen in 500 µl of 0.01% sodium dodecyl sulfate (SDS) and stored at –20°C until ready to process. To assess protein content and algal abundance at each time point, whole anemones were thawed, homogenized for 30 s with an OMNI Tissue Master (OMNI, Kennesaw, GA), and then needle-sheared (with a 25G needle attached to a 1-ml syringe) five times. Following homogenization, protein content was measured using a bicinchoninic acid (BCA) Protein Assay Kit (Thermo Fisher Scientific, Waltham, MA) according to the manufacturer’s protocol, and algal abundance was determined by numerating algal cells remaining in anemones (processed as described by Krediet et al. 2015) using a hemocytometer on an automated cell counter (Corning, Tewksbury, MA). Algal-cell counts were normalized to protein content for each anemone. All measurements were done in triplicate for each sample. The experiment was independently repeated three times, each time with a sample size of 5 anemones per time point for each treatment.

**Figure 2.**
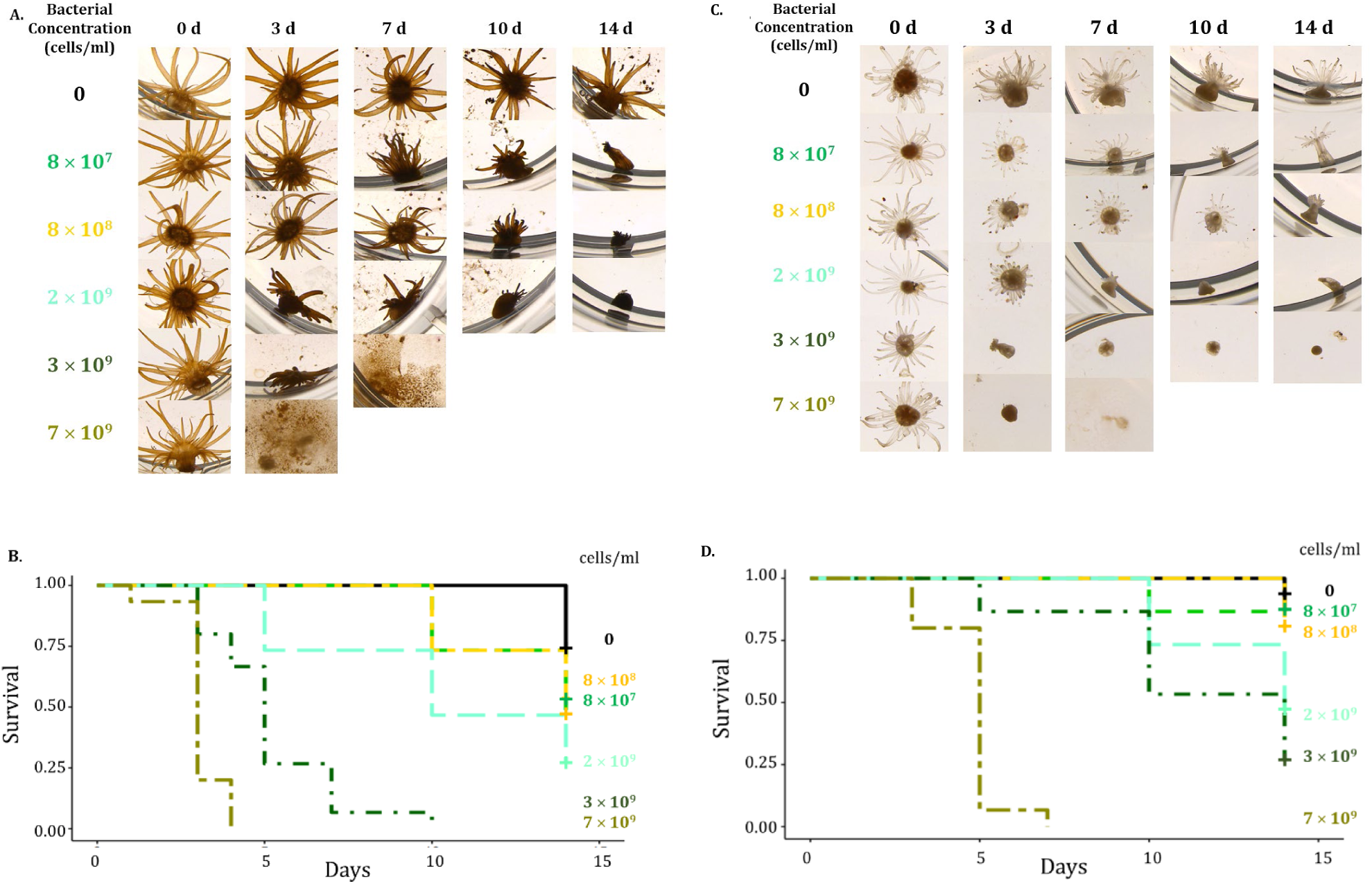
Survival of symbiotic and aposymbiotic Aiptasia over 14 d mostly decrease in response to various inoculation densities of *Vibrio alginolyticus.* Anemones were assessed for signs of disease and death over 14 d following exposure to these *V. alginolyticus* densities: 0, 8 × 10^7^, 8 × 10^8^, 2 × 10^9^, 3 × 10^9^ and 7 × 10^9^ cells ml^-1^. Five anemones were assessed for each density and the experiment was independently repeated three times. **A.** Morphology of representative symbiotic Aiptasia under brightfield microscopy shows tentacle retraction and tissue darkening after 7 d following exposure to all *V. alginolyticus* densities. Tissue degradation began by 14 d at 2 × 10^9^ cells ml^-1^, 5 d at 3 × 10^9^ cells ml^-1^ and 3 d at 7 × 10^9^ cells ml^-1^. **B.** Kaplan-Meier survival curves (χ^2^= 127; *df*= 5; *P*<.001) for symbiotic Aiptasia. Plus signs (+) at the end of curves represent censoring (see Materials and Methods). Survival decreased at all inoculation densities. **C.** Morphology of representative aposymbiotic Aiptasia under brightfield microscopy shows tentacle retraction following exposure to *V. alginolyticus* at inoculation densities greater than 8 × 10^8^ cells ml^-1^ after 3 d. Tissue degradation began by 10 d at 3 × 10^9^ cells ml^-1^ and 7 d at 7 × 10^9^ cells ml^-1^. **D.** Kaplan-Meier survival curves (χ^2^= 107; *df*= 5; *P*<.001) for aposymbiotic Aiptasia. Plus signs (+) at the end of curves represent censoring (see Materials and Methods). Survival decreased at all densities.

### Data analysis

To analyze anemone survival in response to different densities of *R. mobilis*, *A. macleodii*, and *V. alginolyticus*, Kaplan-Meier curves were constructed using anemone status (e.g., live or dead) over 14 d and analyzed with a log-rank test and a Cox proportional-hazards analysis (Rich et al. 2010, Zaragoza et al. 2014, Dey et al. 2020). At the end of each experimental trial, anemones were classified as “censored” if they were alive, and represented on curves as “+” symbols (Rich et al. 2010). The log-rank test compared all curves representing different inoculation densities to each other for each given host type (symbiotic or aposymbiotic) and bacterium (*R. mobilis*, *A. macleodii*, or *V. alginolyticus*). Log-rank test results are shown as a chi-square (χ^2^) and p-value. A Cox proportional-hazards analysis was used to compare changes in host survival between uninoculated and inoculated anemones at each bacterial density, and results are presented as hazards ratios (HR). A hazards ratio is the hazard rate (increase or decrease in survival) of the inoculated group in comparison to the uninoculated control group. When HR=1, there was no change in the hazard (i.e., no change in survival); when HR<1, there was a decreased hazard (i.e., increased survival); and when HR>1, there was an increased hazard (i.e., decreased survival). Survival analyses were conducted using the “survminer” package in R Studio (RStudio, Boston, MA).

To analyze changes in anemone biomass, protein content, and algal over 7 d in response to the same three bacteria, each inoculated at a density of 8 × 10^8^ cells ml^-1^, student’s t-tests were performed in Microsoft Excel to determine any differences between the inoculated and uninoculated anemones at each time point for each of the three health metrics.

## Results

### Aiptasia survival and morphology are influenced by bacterial inoculations and presence of algal symbionts

Survival and morphological assessments were done as a first step in investigating the bacteria’s impact on symbiotic and aposymbiotic Aiptasia. *Ruegeria mobilis* enhanced symbiotic-Aiptasia survival when inoculated at low densities (e.g., 8 × 10^7^ cells ml^-1^) and depressed survival at high densities (e.g., 2 × 10^9^ cells ml^-1^ and greater) (**Figure 1A, B**). Survival of aposymbiotic Aiptasia did not change with 8 × 10^7^, 8 × 10^8^, 2 × 10^9^, and 3 × 10^9^ cells ml^-1^ of *R. mobilis*, and decreased at 7 × 10^9^ cells ml^-1^ over 14 d (**Figure 1C, D**). A log-rank test resulted in significant differences in survival curves of symbiotic (χ^2^= 121; *df*= 5; *P*<.001) and aposymbiotic (χ^2^= 110; *df*= 5; *P*<.001) Aiptasia inoculated with *R. mobilis* at various densities.

Overall, *Vibrio alginolyticus* decreased survival of symbiotic and aposymbiotic Aiptasia over 14 d with little regard to inoculation density. The log-rank test resulted in significant differences in survival of symbiotic (χ^2^= 127; *df*= 5; *P*<.001) and aposymbiotic (χ^2^= 107; *df*= 5; *P*<.001) Aiptasia inoculated with various densities of *V. alginolyticus*. Compared to uninoculated symbiotic hosts, *V. alginolyticus* diminshed survival of symbiotic hosts at all densities (**Figure 2A, B**). *V. alginolyticus* also reduced survival of aposymbiotic hosts, with the exception of 8 × 10^7^ cells ml^-1^ where survival did not change. (**Figure 2C, D**).

*Alteromonas macleodii* decreased symbiotic-Aiptasia survival when inoculated at 2 × 10^9^ cells ml^-1^ or higher, but otherwise did not have an impact on the survival symbiotic anemones at lower densities (**Figure 3A, B**). *A. macleodii* also did not have an impact on aposymbiotic-host survival at the lowest inoculation density (e.g., 8 × 10^7^ cells ml^-1^) but started to diminish anemone survival at 8 × 10^8^ cells ml^-1^ and greater over 14 d (**Figure 3C, D**). A log-rank test resulted in significant differences in survival of symbiotic (χ^2^= 95.9; *df*= 5; *P*<.001) and aposymbiotic (χ^2^= 115; *df*= 5; *P*<.001) Aiptasia inoculated with *A. macleodii* at different densities.

**Figure 3.**
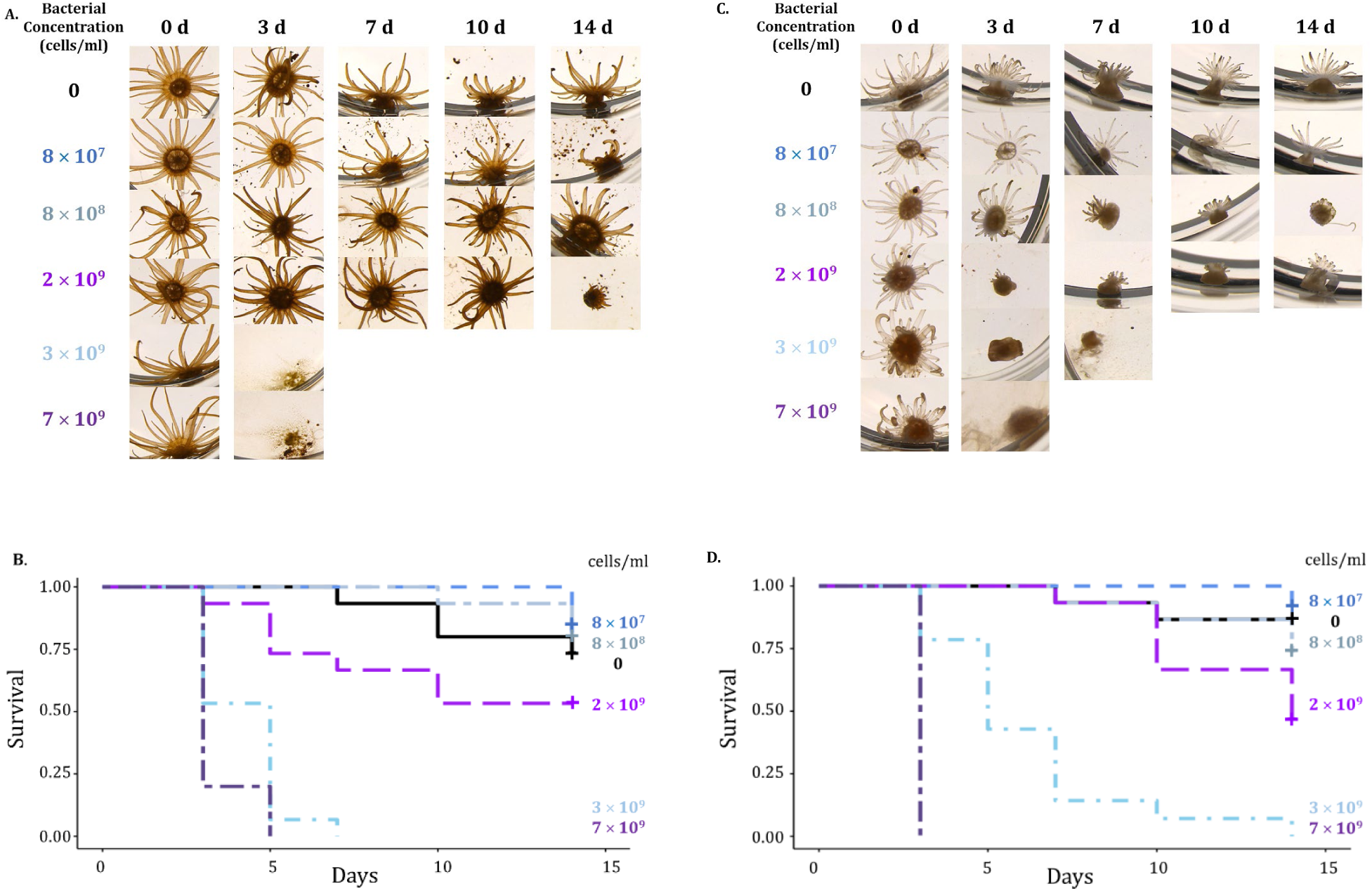
Survival of symbiotic and aposymbiotic Aiptasia over 14 d decrease at higher inoculation densities of *Alteromonas macleodii*. Anemones were assessed for signs of disease and death over 14 d following exposure to these densities of *A. macleodii*: 0, 8 × 10^7^, 8 × 10^8^, 2 × 10^9^, 3 × 10^9^ and 7 × 10^9^ cells ml^-1^. Five anemones were assessed for each density and the experiment was independently repeated three times. **A.** Morphology of representative symbiotic Aiptasia under brightfield microscopy shows little to no tentacle retraction and some host tissue darkening following exposure to *A. macleodii* density of 2 × 10^9^ cells ml^-1^. Tissue degradation began by 14 d at 2 × 10^9^ cells ml^-1^, and 3 d at 3 × 10^9^ cells ml^-1^ and 7 × 10^9^ cells ml^-1^. **B.** Kaplan-Meier survival curves (χ^2^= 95.9; *df*= 5; *P*<.001) for symbiotic Aiptasia. Plus signs (+) at the end of curves represent censoring (see Materials and Methods). Survival decreased at 2 × 10^9^ cells ml^-1^ and greater. **C.** Morphology of representative aposymbiotic Aiptasia under brightfield microscopy shows tentacle retraction at *A. macleodii* densities of 8 × 10^8^ cells ml^-1^ and greater. Tissue degradation began by 3 and 7 d at 7 × 10^9^ cells ml^-1^ and 3 × 10^9^ cells ml^-1^, respectively. **D.** Kaplan-Meier survival curves (χ^2^= 115; *df*= 5; *P*<.001) for aposymbiotic Aiptasia. Plus signs (+) at the end of curves represent censoring (see Materials and Methods). Survival decreased at 8 × 10^8^ cells ml^-1^ and greater.

### High bacterial-inoculation densities are detrimental to Aiptasia survival

The bacterial densities 2 × 10^9^ cells ml^-1^ and greater represent toxic doses to Aiptasia. When symbiotic and aposymbiotic Aiptasia were inoculated with each bacterium at these densities, survival declined compared to uninoculated anemones (**Figures 1-3**). At 2 × 10^9^ cells ml^-1^ for all bacteria, hosts had decreased survival as determined by the Cox proportional-hazards analysis. With the exception of aposymbiotic hosts inoculated with *Ruegeria mobilis*, hazards ratios (HR) were above 1.00, indicating an increased hazard and decreased survival (**Table 1**).

Following inoculation with 2 × 10^9^ cells ml^-1^ of each bacterium, visual signs of health declined in all hosts. All symbiotic anemones showed tissue darkening, while tentacle retraction and tissue degradation occurred in both symbiotic and aposymbiotic hosts by 3 d and animals did not survive past 7 d. For each bacterium, densities of 2 × 10^9^ and 3 × 10^9^ cells ml^-1^ resulted in tissue darkening and tentacle retraction by 7 d and reduced survival in all hosts, with the exception of *R. mobilis* in aposymbiotic Aiptasia, where there was no change in physical appearance or survival. The two lowest densities, 8 × 10^7^ and 8 × 10^8^ cells ml^-1^, gave varying results for each bacterium and host type (**Table 1**). Therefore, the bacterial-inoculation density of 2 × 10^9^ cells ml^-1^ appears to be a threshold for Aiptasia survival.

### Aiptasia biomass, protein content, and algal abundance are impacted by bacterial inoculations and presence of algal symbionts

Three different metrics—biomass, protein content, and algal abundance—were adopted to assess other dimensions of anemone health in addition to survival, following inoculation with 8 × 10^8^ cells ml^-1^ of each bacterium. Compared to uninoculated Aiptasia, symbiotic anemone biomass increased by 8% at 7 d (*P*=.01) when inoculated with *R. mobilis*, whereas host biomass was not impacted by *A. macleodii* but decreased by 30% at 2 d (*P*<.001) and 27% by 7 d (*P*<.001) when inoculated with *V. alginolyticus* (**Figure 4A**). Among aposymbiotic Aiptasia, biomass increased at 1 d by 11% (*P*<.01) following inoculation with *R. mobilis*, but no significant impact occurred at other experimental time points (**Figure 4B**). Biomass decreased at 2 d by 8% (*P*<.001) following inoculation with *A. macleodii*, but had no impact by 7 d (**Figure 4B**). Finally, following inoculation with *V. alginolyticus*, biomass decreased by 36% at 3 d (P<.001) and continued to decrease all the way by 54% at 7 d (*P*<.01) relative to uninoculated anemones (**Figure 4B**).

**Figure 4.**
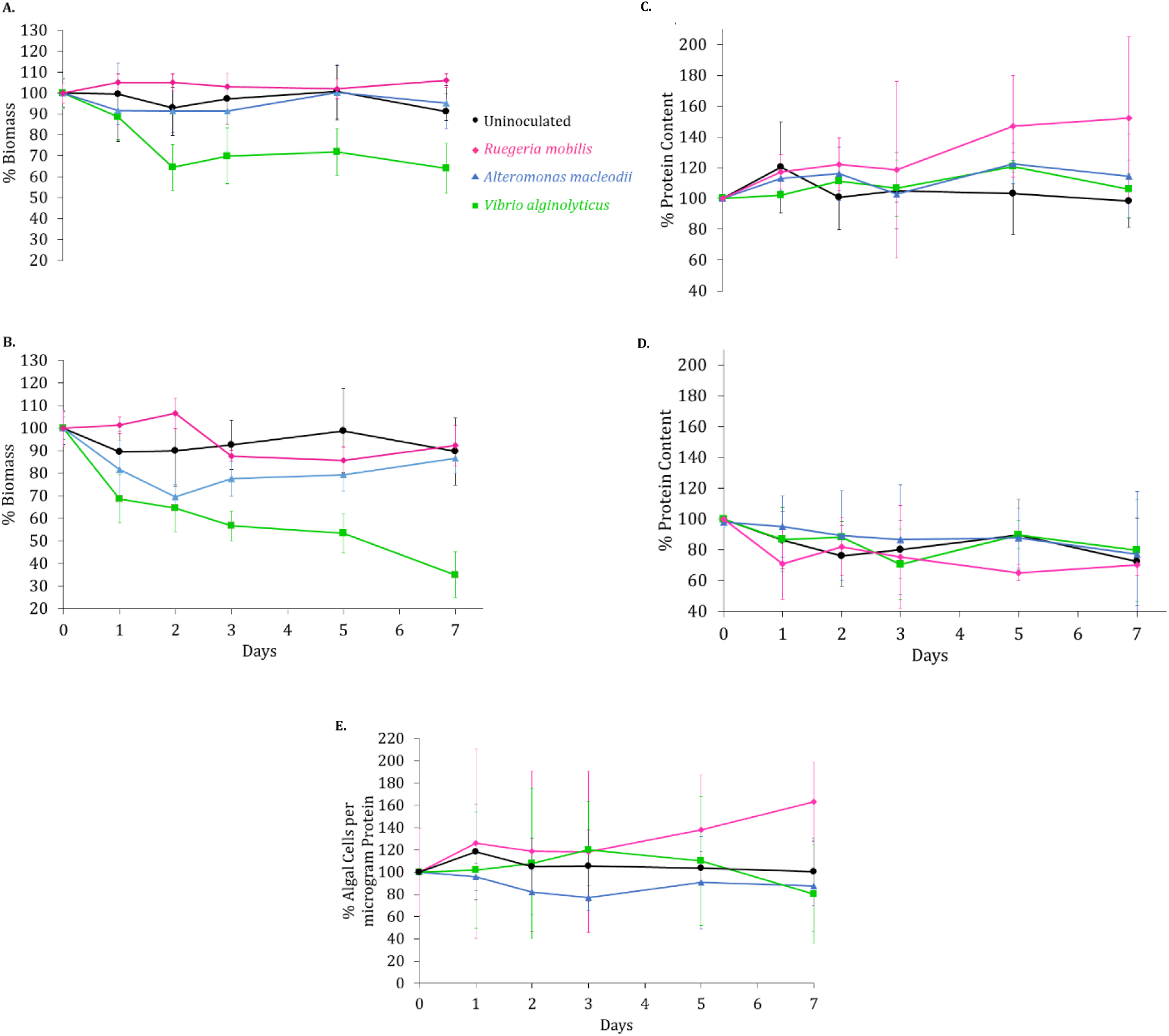
*Ruegeria mobilis* enhances biomass, protein content, and algal abundance of Aiptasia over 7 **d.** Symbiotic and aposymbiotic Aiptasia were independently inoculated with *Ruegeria mobilis, Vibrio alginolyticus, and Alteromonas macleodii* at 8 × 10^8^ cells ml^-1^. Five anemones were assessed for each bacterium on each day and the experiment was independently repeated three times. **A.** In symbiotic hosts, when compared to uninoculated hosts, *R. mobilis* increases host biomass by 8% at 7 d (*P*=.01), *A. macleodii* does not impact biomass, and *V. alginolyticus* decreases host biomass by 30% at 2 d (*P*<.001) and 27% at 7 d (*P*<.001). **B.** In aposymbiotic anemones, *R. mobilis* increases anemone biomass at 1 d by 11% (*P*<.001) and has no impact for the remainder of the experiment. *A macleodii* decreases biomass at 2 d by 8% (*P*<.001) and has no impact by 7 d. *V. alginolyticus* decreases host biomass by 36% at 3 d (*P*<.001) and 54% at 7 d (*P*<.01). **C.** In symbiotic anemones, *R. mobilis* increases host protein content by 43% at 5 d (*P*<.001) and 50% at 7 d (*P*<.001), while *V. alginolyticus* and *A. macleodii* have no impact. **D.** In aposymbiotic anemones, *R. mobilis* decreases host protein content by 24% at 5 d (*P*=.009) but does not impact protein content at the end of 7 d. *A. macleodii* and *V. alginolyticus* do not impact host-protein content. **E.** Algal abundance within symbiotic Aiptasia. Percentage of algal cells was normalized to host-protein content. At 7 d, *R. mobilis* increases algal abundance by 63% (*P*=.02), while *V. alginolyticus* decreases algal abundance by 20% (*P*=.05). *A. macleodii* has no impact on algal abundance.

Protein content of symbiotic hosts increased with inoculation of *R. mobilis* and decreased with inoculation of *V. alginolyticus* (**Figure 4C**), while protein content of aposymbiotic hosts was unchanged by 7 d across all treatments. In symbiotic Aiptasia, host-protein content increased by 43% (*P*<.001) at 5 d and 50% (*P*<.001) by 7 d following *R. mobilis* inoculation (**Figure 4C**). Following inoculation with *V. alginolyticus* and *A. macleodii* host-protein content did not change (**Figure 4C**). In aposymbiotic Aiptasia, *R. mobilis* reduced host-protein content by 24% at 5 d (*P*=.009) but did not impact protein content at the end of 7 d, and *A. macleodii* and *V. alginolyticus* did not impact host protein content altogether (**Figure 4D**).

Both *R. mobilis* and *V. alginolyticus* appear to have an impact on host algal abundance (i.e., the number of algal cells remaining in the host over time). Algal abundance increased at 7 d by 63% (*P*=.02) following inoculation with *R. mobilis*, while algal abundance decreased by 20% at 7 d (*P*=.05) following inoculation with *V. alginolyticus* (**Figure 4E**)*. A. macleodii* had no impact on algal abundance at all (**Figure 4E**). Assessments of survival, morphology, biomass, protein content, and algal abundance of anemones are summarized to determine the impacts on the three bacterial species on host health (**Table 2**). It was predicted and, in the end, observed that *R. mobilis* had an overall positive effect on the host, *V. alginolyticus* had an overall negative effect, and *A. macleodii* had no apparent effect on the host (**Table 2**).

**Table 2.**
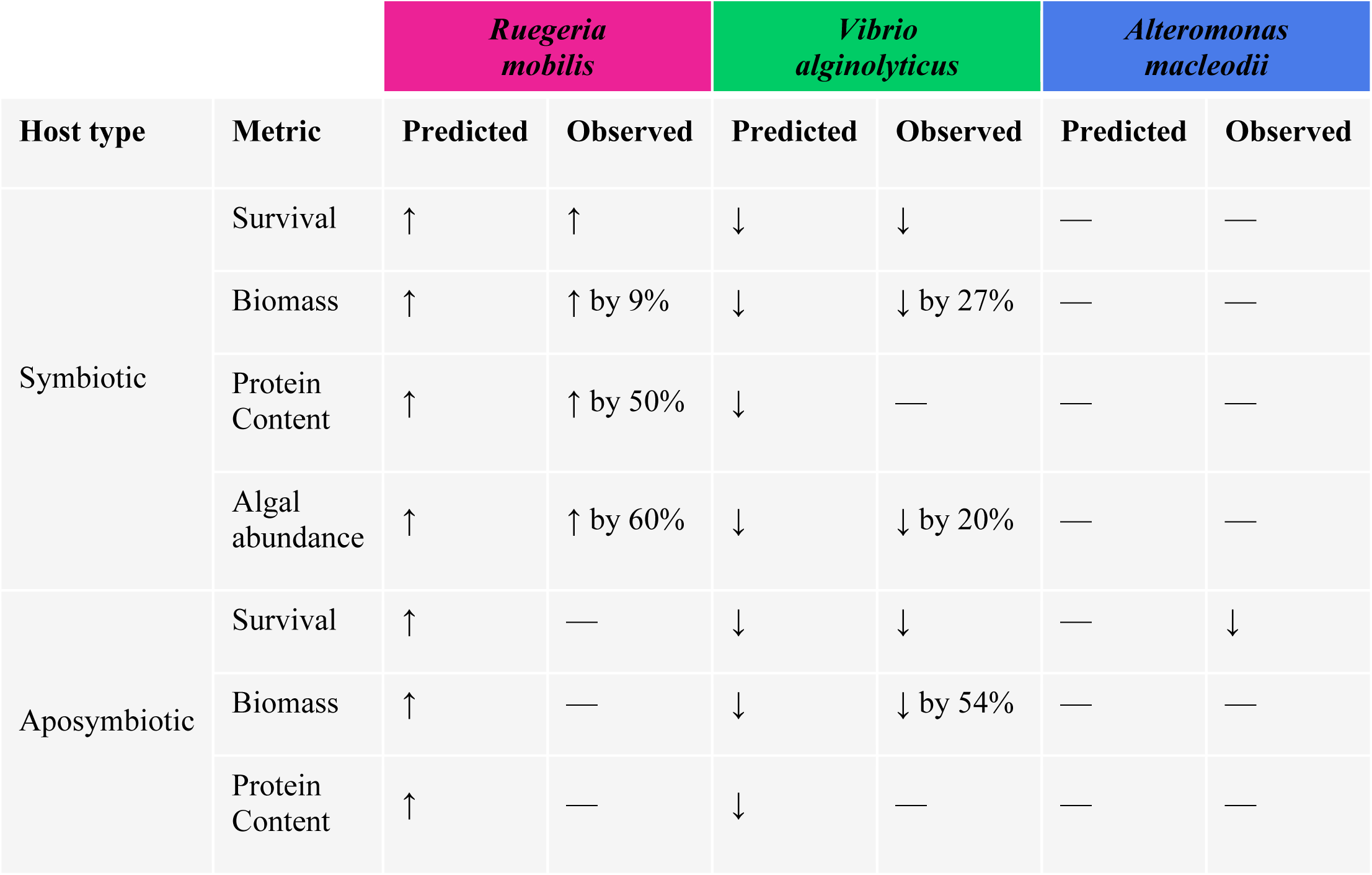
Summary of predicted and observed outcomes on Aiptasia following exposure to three bacterial species, *Ruegeria mobilis*, *Vibrio alginolyticus*, and *Alteromonas macleodii*. Survival, biomass, protein content, and algal abundance were used to determine impact on symbiotic and aposymbiotic hosts. Impact is indicated by arrows, where “↑” indicates increase, “↓” indicates decrease, and “—” indicates no change in the metric following exposure to each bacterial species.

|  |  | *Ruegeria mobilis* |  | *Vibrio alginolyticus* |  | *Alteromonas macleodii* |  |
| --- | --- | --- | --- | --- | --- | --- | --- |
| Host type | Metric | Predicted | Observed | Predicted | Observed | Predicted | Observed |
| Symbiotic | Survival | ↑ | ↑ | ↓ | ↓ | — | — |
|  | Biomass | ↑ | ↑ by 9% | ↓ | ↓ by 27% | — | — |
|  | Protein Content | ↑ | ↑ by 50% | ↓ | — | — | — |
|  | Algal abundance | ↑ | ↑ by 60% | ↓ | ↓ by 20% | — | — |
| Aposymbiotic | Survival | ↑ | — | ↓ | ↓ | — | ↓ |
|  | Biomass | ↑ | — | ↓ | ↓ by 54% | — | — |
|  | Protein Content | ↑ | — | ↓ | — | — | — |

## Discussion

As ocean temperatures are rising, the cnidarian microbial community changes, negatively impacting holobiont health (Yellowlees et al. 2008; Littman et al. 2010; Rädecker et al. 2021). Bacteria associated with cnidarians may confer some advantage for managing heat stress (Peixoto et al. 2017; Rosado et al. 2019). Inoculating corals with beneficial microorganisms may, therefore, give corals an advantage. The specific physiological impacts of individual bacterial species on the holobiont are unknown and, in this study, we investigated three bacterial species (*Ruegeria mobilis*, *Vibrio alginolyticus*, and *Alteromonas macleodii*) associated with corals and the sea anemone Aiptasia to determine their impacts on cnidarian-host health. Overall, anemones exhibited distinct responses to different bacterial species; furthermore, bacterial-inoculation densities and the symbiotic state of the anemones (i.e., whether they contain algal symbionts or not) were additional factors that determined the varying levels of impact of each bacterium.

### Three different marine bacteria with distinct impacts on cnidarian-host health

*Alteromonas* spp. are ecologically important to marine ecosystems, yet their functional role in the cnidarian holobiont required further investigation. A limited number of studies have pointed to their role in the degradation of reactive oxygen species (ROS) (Hernandez-Agreda et al. 2018). In marine environments, *Alteromonas* spp. dominate the microhabitat created by algal blooms, where they degrade algal polysaccharides and remove ROS. By the same token when these bacteria are inside of a host, they have the potential capacity to remobilize carbon and protect hosts from hydrogen-peroxide damage (Morris et al. 2011; Nuemann et al. 2015; Koch et al. 2019). While ROS activity was not measured in the present study, *A. macleodii* had little to no impact on Aiptasia health (**Table 2**), as it only decreased anemone survival at high bacterial densities (**Figure 3**) but did not change biomass, protein content, or algal abundance at all (**Figure 4**). Though *A. macleodii* is very abundant in laboratory strains of Aiptasia (Binsarhan 2015, Hartman et al. 2020, Sydnor et al. 2023), it does not appear to contribute much to the host.

Although the *Alteromonas* genus is also prevalently associated with corals, Allers and colleagues (2008) suggest that its main action is not any contribution to the holobiont, but to transfer organic carbon to the outer food web. *A. macleodii* produces particulate organic carbon and particulate nitrogen at high levels within the coral mucus (Allers et al. 2008). When this mucus is released by corals, this bacterial particulate matter is consumed by organisms at higher trophic levels, such as phytoplankton. Taken together with our results, *A. macleodii* does not appear to have a central role in the maintenance of host health, or that was not assessable by the metrics used in this study.

Some members of the *Vibrio* genus have long been associated with common coral diseases. *Vibrio* spp. are five times more abundant in unhealthy than in healthy corals (Hernandez-Agreda et al. 2017; Rubio-Portillo et al. 2018). When temperatures increase, *Vibrio* spp. can overgrow and dominate the coral microbiome, while becoming pathogenic in the process, leading to coral disease (Bourne et al. 2008; Mao-Jones et al. 2010). Though the exact pathogenic pathways have not been established in *V. alginolyticus* specifically, there are virulence genes that encode for hemolysins and proteases (Hernández-Robles et al. 2016). Furthermore, *V. alginolyticus* forms capsules to protect itself from phagocytosis by host cells and adhere to surfaces for biofilm formation (Hernández-Robles et al. 2016), allowing its persistence within or around the host.

*Vibrio* spp. are also commonly associated with Aiptasia and, most notably, their abundance dramatically increases in heat-stressed, aposymbiotic animals (Sydnor et al. 2023). The most dominant species identified in that study were *V. harveyi*, *V. parahemolyticus*, and *V. alginolyticus*. Other *Vibrio* spp., such as *V. coralliilyticus* and *V. shiloi*, also cause disease and tissue disintegration in Aiptasia (Zaragoza et al. 2020). In the present study, we specifically isolated *V. alginolyticus* from Aiptasia and found that it impaired host health (**Table 2**). *V. alginolyticus* diminished host survival and biomass of both symbiotic and aposymbiotic Aiptasia (**Figures 2, 4**), as well as protein content and algal abundance of symbiotic Aiptasia (**Figure 4**).

Despite no loss in algal symbionts within Aiptasia in response to *V. coralliilyticus* or *V. shiloi* (Zaragoza et al. 2020), *Vibrio* spp. have been implicated in bacterial-induced bleaching of corals (Rosenberg et al. 2004). Our data support the hypothesis that bacteria are an important factor in coral bleaching, with stress and subsequent Symbiodiniaceae expulsion induced by bacterial infections (Rosenberg et al. 2004; Zhou et al 2020).

Furthermore, our results also support previous findings of darkened cnidarian tissue as an indicator of a host-immune response to bacterial infections (Palmer et al. 2011; Zaragoza et al. 2020). In this study, tissue darkening of symbiotic hosts was elicited by all *V. alginolyticus* inoculation densities (**Figure 2**). This is likened to the production of melanin as a result of phenoloxidase activity in one pathway of innate immunity (Palmer et al. 2011; Zaragoza et al 2020). Melanin is produced to kill invading microorganisms, scavenge oxygen radicals released from Symbiodiniaceae during stress, and absorb light during bleaching (Palmer et al. 2011). In corals, peroxidase activity and melanin production both increase near disease lesions (Palmer et al. 2011). Further investigation into the mechanisms underlying melanin production in Aiptasia in response to pathogenic infections is clearly warranted.

Aiptasia may harbor beneficial microbes that have the capacity to outcompete pathogens or enhance their survival, but to date, no specific bacterial symbiont(s) have been definitively identified for this model organism. Rhodobacteraceae was one of the two most abundant bacterial families consistently maintained with Aiptasia despite various environmental treatments (Sydnor et al. 2023), and one of the strains we isolated and tested, *R. mobilis*, belonged to this family. *Ruegeria* spp. release tropodithietic acid and dimethyl sulfide, both antibacterial compounds that have the capacity to inhibit algal and coral pathogens (Raina et al. 2009; Sonnenschein et al. 2016) and may, thus, protect both the cnidarian host and its algal symbionts.

In addition, *Ruegeria* spp. are catalase– and oxidase-positive, suggesting their ability to scavenge ROS (Wirth and Witman 2019). This function may potentially protect corals from bleaching and death, as excess amounts of ROS released by Symbiodiniaceae within the host under thermal stress are potential causes of symbiosis breakdown and host-tissue damage (Wietheger et al. 2018). Bacteria, like *Ruegeria* spp., may be able to reduce this oxidative damage and potentially counter coral bleaching, though more studies need to be done to fully understand their role (Dungan et al. 2021).

In the present study, we observed many benefits of *R. mobilis* in improving host survival (**Figure 1**), biomass, protein content, and algal abundance (**Figure 4**), but only in symbiotic Aiptasia with no apparent impact on aposymbiotic animals (**Table 2**). Thus, the presence of Symbiodiniaceae seems to matter and suggests that *R. mobilis* might have a significant impact on the host through these algal symbionts. It is unclear what specific interactions between the two may be, or the mechanisms by which these interactions may in turn benefit the host. However, their association is not novel or surprising. *Ruegeria* spp. are commonly associated with dinoflagellate algae, found in Symbiodiniaceae cultures across the globe (Sonnenschein et al 2017; Matthews et al. 2020). Class-specific probes for fluorescence *in situ* hybridization identified Alphaproteobacteria (which includes *Ruegeria* spp.) to be associated with five different Symbiodiniaceae species originally isolated from corals and Aiptasia (Maire et al. 2021). When bacteria from 11 Symbiodiniaceae cultures were separated with sterile washes into “loosely” (extracellular to the host and not attached), “closely” (tightly attached to the outer surface), and “intracellular” fractions, metabarcoding revealed that the Rhodobacteraceae family (which includes *R. mobilis*) was found to be in all three Symbiodiniaceae fractions (Maire et al. 2021). Future directions may consider advanced microscopy and other means to specifically elucidate the interactions between *R. mobilis* and Symbiodiniaceae, both *in hospite* and *ex hospite*.

### Healthy vs. bleached cnidarians may respond differently to probiotics and pathogens

The presence of algal symbionts within the host may influence host-bacterial interactions, and possibly bacterial colonization and growth. Aiptasia responded differently to each bacterial species depending on the presence of Symbiodiniaceae within anemones (**Table 2**), suggesting that algae play some role in how bacteria interact with the cnidarian holobiont. This was not only apparent with *R. mobilis*, as mentioned, but *V. alginolyticus* across all densities tested also decreased symbiotic-anemone survival to a greater extent than aposymbiotic-host survival (**Figure 2**). While we have only observed this in the few bacterial species tested in this study, algal symbionts are likely providing necessary nutrients and an expanded niche through the phycosphere to support a large number and diversity of bacteria (Frost et al. 2008; Garrido et al. 2020; Costa et al. 2021). As bacteria are found extra– and intracellularly within Symbiodiniaceae cells (Matthews et al. 2020; Maire et al. 2021; Marangon et al 2021), resource exchange between the two types of microbes provides stability and resilience to the coral holobiont (Matthews et al. 2020). Without algal symbionts present, bacteria may be carbon-limited (Frost et al 2005; Frost et al. 2008) and not thrive within the host or contribute to its health as much.

The differential responses to bacterial inoculations exhibited by symbiotic and aposymbiotic anemones should be considered when assessing the efficacy of probiotic administration to corals as a proposed form of coral conservation (Rosado et al. 2019; Blackall et al. 2020). While microbiome manipulation has been gaining traction due to promising results that include improving host physiology and survival, mitigating host bleaching, augmenting pathogen defense, enhancing calcification rates, and degrading oil or minimizing oil-spill effects on corals (Rosado et al. 2019; Silva et al. 2021; Zhang et al. 2021), our results in a cnidarian laboratory model indicate distinct differences between hosts with and without their algal symbionts. This may imply that corals that have already bleached may respond very differently to probiotics than healthy corals with their dinoflagellate-symbiosis intact. For example, *R. mobilis* may be considered as a possible probiotic with its ROS-scavenging capabilities, but may not confer an advantage to fully bleached corals (that have lost all their algal symbionts). Thus, probiotics are best used as a preventative, not reactive, measure to coral bleaching.

When evaluating the use of beneficial bacteria for microbiome manipulation, another factor to consider is bacterial density. Our results demonstrate that high bacterial densities weaken host health, regardless of the bacterial species inoculated, and when utilizing bacteria to improve cnidarian health, densities below 2 × 10^9^ cells ml^-1^ may be a helpful guideline. Too many bacterial cells in the seawater may deplete the host of oxygen, while a high density of any single bacterium could disrupt the balance of the microbiome, with inoculated bacteria outcompeting other bacteria that maintain metabolic homeostasis within the holobiont (Kline et al. 2006). Moreover, the bacterial carrying capacity of Aiptasia is around 10^5^ cells polyp^-1^, similar that of corals of about 10^6^ cells cm^-2^, with the host unable to support higher bacterial densities (Costa et al. 2021). Inoculations with lower bacterial densities may be required to improve host health. Bacterial densities of 10^6^ and 10^7^ cells ml^-1^ were used to inoculate corals with beneficial microbes and health was improved (Damjanovic et al. 2019; Rosado et al. 2019); in this study, 10^7^ and 10^8^ cells ml^-1^ enhanced Aiptasia health. Therefore, future studies should focus on determining how many of these inoculated bacterial cells are physically interacting (and potentially colonizing) the host.

Coral reefs are bleaching and dying at an unprecedented rate due to global climate change and anthropogenic disturbances. Coral-reef-conservation techniques, such microbiome manipulation, must be further investigated and implemented to mitigate coral destruction. Before microbiome manipulation can be successfully utilized *in situ* in marine environments, we must fully understand the impacts individual bacterial species have on the cnidarian holobiont. This study clearly highlights the beneficial impacts of one key bacterial species, *R. mobilis*, on the cnidarian model Aiptasia. Furthermore, this study has demonstrated that Aiptasia can be a powerful model for investigating coral-bacterial diseases. *V. alginolyticus* is a known coral pathogen but, prior to this study, has never been observed to cause disease in Aiptasia. Though we were able to observe specific impacts of these bacterial species on the host under ambient environmental conditions, performing these experiments at increased temperatures will determine whether a beneficial bacterium like *R. mobilis* can truly protect corals from heat stress.

## Acknowledgements

The authors would like to thank Emily Fleming, Kris Blee, and members of the Tran Laboratory research group for helpful discussions about this work. Some content of this manuscript previously appeared online in a different form as part of KR’s Master’s thesis (Romo, 2021). This work was supported by a Student Award in Research and Creativity to KR, along with a CSU STEM-NET grant to CT.

